# Cortical morphology predicts placebo response in multiple sclerosis

**DOI:** 10.1101/825638

**Authors:** Mariya V. Cherkasova, Jessie F. Fu, Michael Jarrett, Poljanka Johnson, Shawna Abel, Roger Tam, Alexander Rauscher, Vesna Sossi, Shannon Kolind, David Li, A. Dessa Sadovnick, Lindsay Machan, J. Marc Girard, Francois Emond, Reza Vosoughi, Anthony Traboulsee, A. Jon Stoessl

## Abstract

Although significant insights have been gained into the neural mechanisms of acute placebo responses, less is known about the mechanisms of longer-term placebo responses, such as those seen in clinical trials, or the interactions between these mechanisms and brain disease. We examined neuropathological and morphological brain correlates of placebo responses in a randomized clinical trial of a controversial endovascular treatment (“liberation therapy”) for multiple sclerosis. Patients were randomized to receive either balloon or sham extracranial venoplasty and followed for 48 weeks. The trial did not support therapeutic efficacy of venoplasty, but a subset of both venoplasty- and sham-treated patients reported an improvement in health-related quality of life that peaked at 12 weeks following treatment, suggesting a placebo response. Placebo responders had higher lesion activity than placebo non-responders. Although placebo responders did not differ from non-responders in terms of total normalized brain volume, regional grey or white matter volume or cortical thickness, graph theoretical analysis of cortical thickness covariance showed that placebo non-responders had a more homogenous cortical thickness topology with a more small-world-like architecture. In placebo non-responders, lesion load inversely predicted cortical thickness in primary somatosensory and motor areas, association areas, precuneus and insula, primarily in the right hemisphere. In placebo responders, lesion load was unrelated to cortical thickness. The neuropathological process in MS may result in a cortical configuration that is less suited to functional integration and less capable of generating a sustained placebo response.

## INTRODUCTION

Current understanding of the neurobiology of placebo effects comes primarily from laboratory studies of acute placebo interventions. Longer term placebo responses, such as those in clinical trials, have been less studied but appear to rely on structural and functional brain connectivity (Tétreault et al., 2016)(Hashmi et al., 2012)(Vachon-Presseau et al., 2018)(Liu et al., 2017) and involve modulation of fMRI-derived (Wanigasekera et al., 2018) and metabolic networks (Mayberg et al., 2002)(Ko et al., 2014)(Niethammer et al., 2018).

Unlike acute laboratory placebo responses, often studied in healthy participants, those of patients with chronic conditions in clinical trials and real-world settings may reflect a yearning for improvement, tempered by varying levels of hope, prior therapeutic experiences and acceptance of risk for a chance at recovery. In neurobehavioural disorders, placebo mechanisms may interact with neuropathological processes. For example, Alzheimer’s patients show a reduced capacity for placebo analgesia, which has been linked to disrupted connectivity of the prefrontal cortex with the rest of the brain (Benedetti et al., 2006). However, little is known about the interactions between placebo responses and brain disease.

To address this question, we examined neuropathological and structural neural correlates of placebo responses of multiple sclerosis (MS) patients undergoing a randomized clinical trial (RCT) of a controversial extracranial venoplasty procedure dubbed the “liberation therapy”. The treatment was based on the now discredited notion that chronic cerebrospinal venous insufficiency contributes (CCSVI) to MS pathogenesis (Traboulsee et al., 2013). Owing to the initially promising results of uncontrolled, unblinded studies (Dake, Dantzker, Bennett, & Cooke, 2012)(Hubbard et al., 2012)(Zamboni et al., 2009)(Radak et al., 2014)(Salvi, Buccellato, & Galeotti, 2012)(Zagaglia, Balestrini, & Perticaroli, 2013) and the associated publicity, many patients viewed it as a potential cure and sought it out despite the potential risks and the skepticism of the scientific community. However, venoplasty proved ineffective in two double-blind sham-controlled RCTs, one by the pioneers of the procedure (Zamboni et al., 2018), and the other by our group (Traboulsee et al., 2018). In the latter, while venoplasty was not superior to sham venoplasty on any outcome measure, a subset of both venoplasty- and sham-treated patients experienced a significant transient improvement in self-reported health-related quality of life suggesting a placebo response (Figure 1A). This presented a unique opportunity to examine the neural mechanisms of placebo responses in a real-world context.

**Figure 1:**
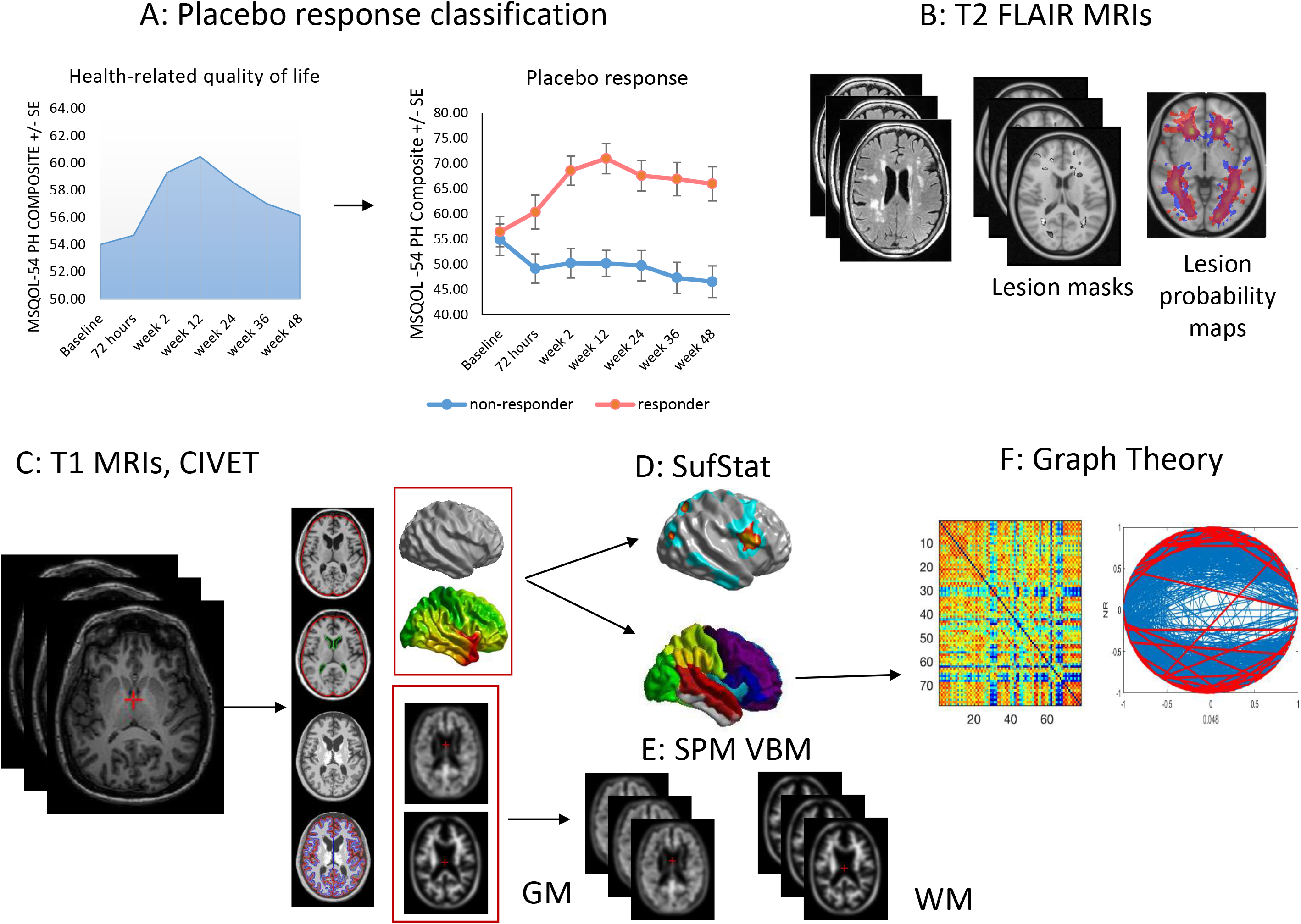
Method overview.

We examined MRI-based predictors of this placebo response (Figure 1). These included lesion load, activity and location, global and regional brain volume and cortical thickness. Because brain disease can disrupt placebo response, we hypothesized that placebo non-responders would have increased white matter lesions as well as grey matter atrophy. Based on the domain-general nature of this placebo response (i.e. health-related quality of life), we expected these increases to be most prominent in regions broadly implicated in reward expectancy and interception such as the prefrontal cortex, striatum and insula. Besides standard morphometric analyses, we performed a graph theoretical analysis of cortical thickness (CT) covariance to characterize its inter-regional relationships. Graph theory is a modality invariant framework that represents complex systems as networks and describes their organization using a set of common metrics. In graph theoretical terms, the brain is viewed as a network of regions (“nodes”) connected via links (“edges”) representing white matter tracts, structural covariance or functional connections. While the neurobiological significance of structural covariance networks and more specifically CT networks is not entirely clear, CT is known to covary between structurally and functionally connected regions (Alexander-Bloch, Giedd and Bullmore, 2013).

This covariation appears to reflect stronger synaptic connectivity between those regions that are microstructurally similar (Suarez, Markello, Betzel, & Misic, 2020)(Seidlitz et al., 2018). Previous studies applying graph theory to CT covariance in MS have found increased network segregation and enhancement of local properties in early disease (Fleischer et al., 2019)(Muthuraman et al., 2016) with a shift in both local and global properties towards more “regular” or uniform networks with advancing disease (Tewarie et al., 2014)(He et al., 2009). Studies of diffusion tensor imaging (DTI) based structural networks (Fleischer et al., 2017)(Shu et al., 2011) and functional connectivity (Tewarie et al., 2014) yielded convergent findings. We hypothesized that CT covariance networks would be more anomalous in placebo non-responders. We specifically focused on three key graph metrics of network segregation and integration: clustering coefficient - a measure of segregation; pathlength – a measure of integration; and the small-world index – a derivative measure describing overall network topology.

## MATERIALS AND METHODS

### Participants

Participants with relapsing remitting (RRMS), secondary (SPMS) and primary progressive (PPMS) MS were recruited between May 29, 2013 and Aug 19, 2015 from four Canadian academic centers: 1) University of British Columbia Hospital, Vancouver; 2) Health Sciences Centre, Winnipeg; 3) CHUM, Hôpital Notre-Dame, Montreal; 4) Hôpital Enfant-Jesus, Québec. Inclusion criteria were: age 18-65 years, diagnosis of definite MS by the 2010 McDonald criteria (Polman et al., 2011), an Expanded Disability Status Score (EDSS) (Kurtzke, 1983) between 0 (i.e. minimal disability) and 6.5 (i.e. using bilateral aids to walk), neurologically stable disease within the 30 days prior to screening, and fulfillment of at least two ultrasound criteria for CCSVI see (Traboulsee et al., 2018) for a detailed description of the trial’s methods and entry criteria. Participants on standard disease-modifying therapies were permitted to continue on the medication, and changes were allowed for on study relapses after randomization. Exclusion criteria were treatment with vasodilators, parasympathomimetics, sympathicolytics, calcium channel blockers, previous venoplasty and/or stenting, previous jugular or subclavian central line or major neck surgery or radiation, previous contrast allergy, inability to undergo MRI, and inadequate medical records confirming diagnosis and disease course. The clinical research ethics boards at the four participating centers approved the study protocol, and participants gave written informed consent. Of the total 104 MS participants, we analyzed the data of 88 who had T1-weighted MRIs of sufficient quality for CT analyses. For the remaining scans, signal intensity at the lateral extremes was too low for successful surface extraction.

Although the primary purpose of this study was to compare the placebo responders to non-responders in the trial, we used MRIs from 43 gender and age-matched healthy controls (30 females, age: 52.98 ± 8.93) to provide a benchmark for graph theory analysis of placebo responders vs. non-responders to help determine which CT pattern was more normative. Six of these scans were acquired at Site 1; the rest were obtained from the Open Access Series of Imaging Studies repository (OASIS-3, www.oasis-brains.org, RRID:SCR_007385).

### Experimental Design and Procedure

Eligible participants were randomized 1:1 to either sham or active balloon venoplasty of all narrowed veins under study. Stratified randomization (RRMS versus progressive MS course) at each site was completed by a permuted-block size of six. Venography was performed under conscious sedation and the duration of time within the angiography suite was uniform for both venoplasty and sham treated participants. A 5-French diagnostic catheter was introduced through the common femoral vein to selectively catheterize the right and left internal jugular veins as well as the azygos vein. The venoplasty participants were treated with an angioplasty balloon 2mm greater than the nominal vein diameter which was inflated for 60 seconds. The participants randomized to sham had a catheter that was advanced across the stenosis and left for 60 seconds.

After randomization and intervention, participants were followed for 48 weeks with MRI, ultrasound, clinical assessments and patient-reported outcome scales including Multiple Sclerosis Quality of Life −54 (MSQOL-54) (Vickrey et al., 1995). For MRI, T1 weighted images with and without gadolinium enhancement, T2 weighted images and fluid attenuated inversion recovery (FLAIR) images were obtained. MRI acquisition parameters are given in Table 1.

**Table 1.**
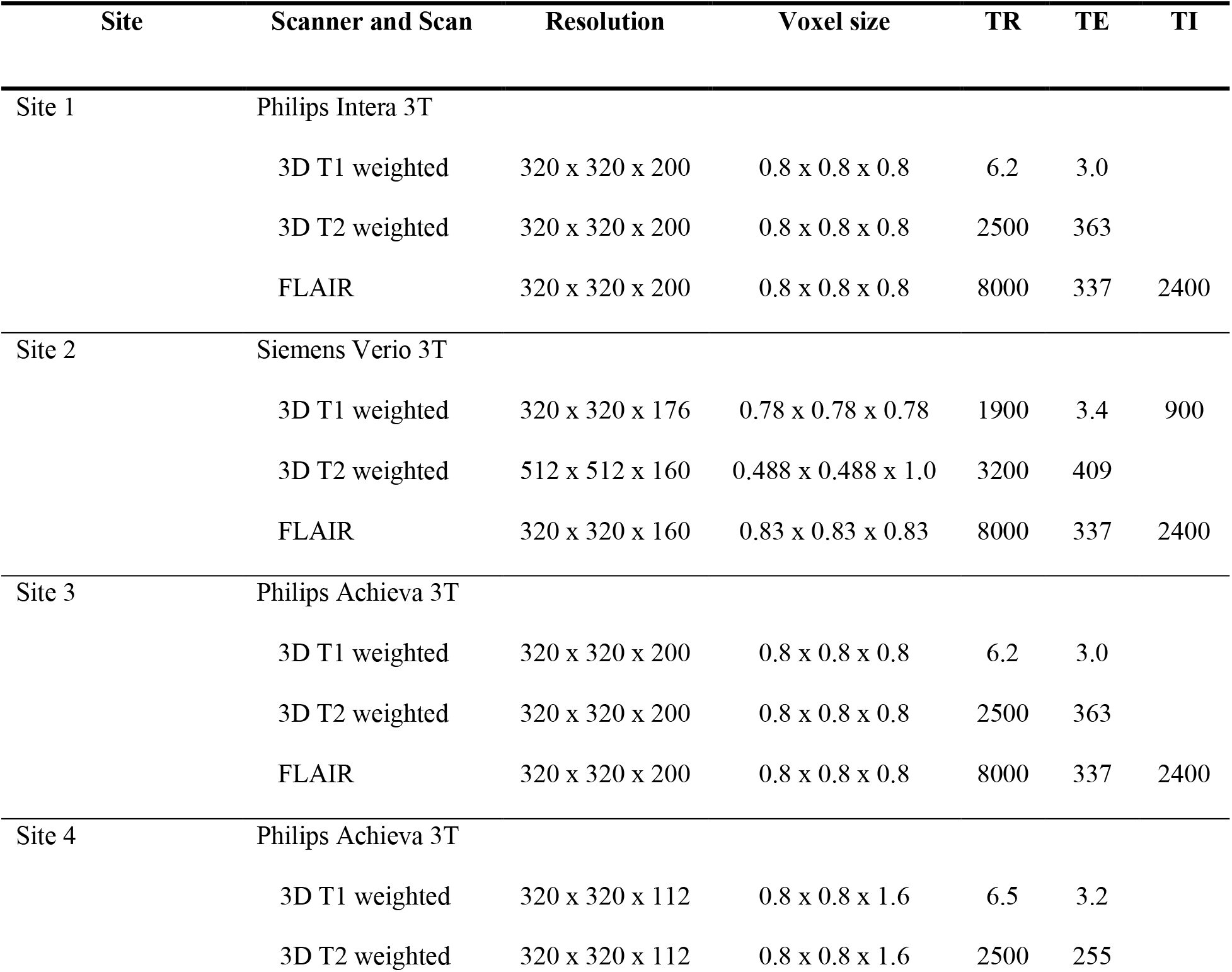
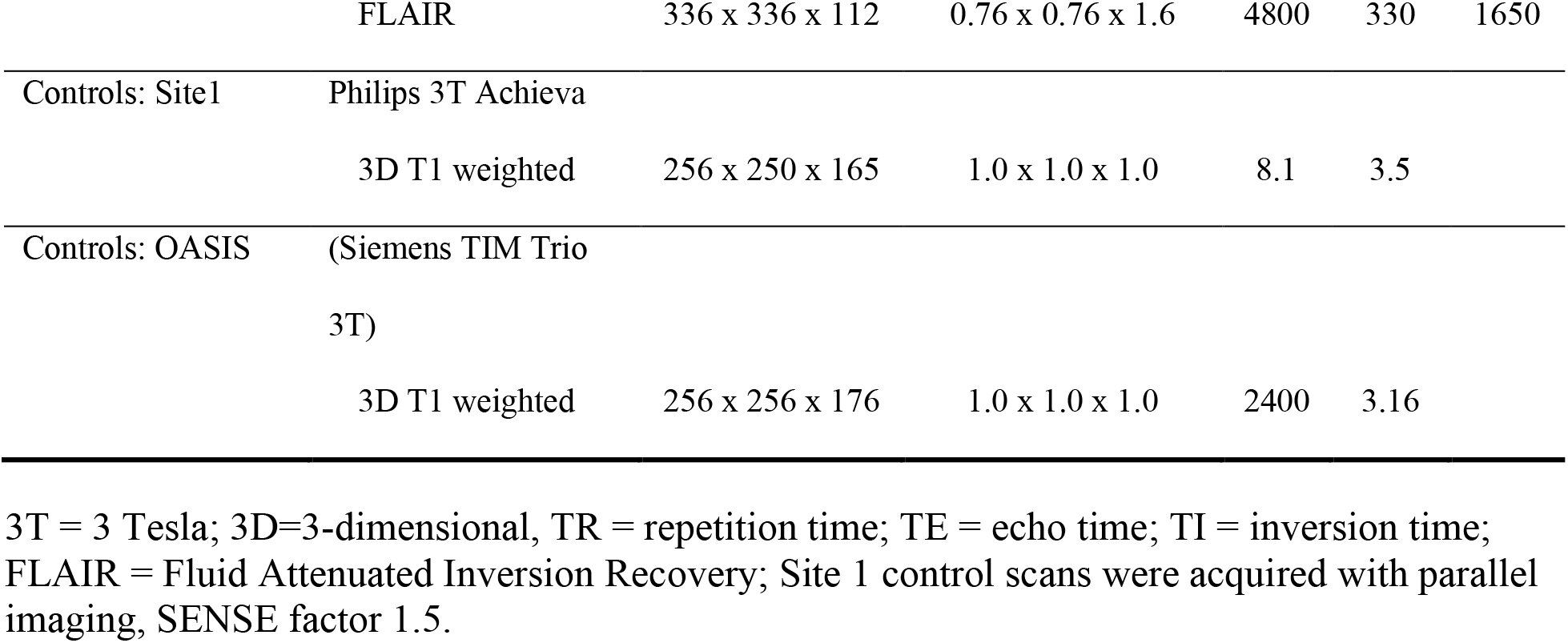
MRI acquisition parameters

| Site | Scanner and Scan | Resolution | Voxel size | TR | TE | TI |
| --- | --- | --- | --- | --- | --- | --- |
| Site 1 | Philips Intera 3T |  |  |  |  |  |
|  | 3D T1 weighted | 320 x 320 x 200 | 0.8 x 0.8 x 0.8 | 6.2 | 3.0 |  |
|  | 3D T2 weighted | 320 x 320 x 200 | 0.8 x 0.8 x 0.8 | 2500 | 363 |  |
|  | FLAIR | 320 x 320 x 200 | 0.8 x 0.8 x 0.8 | 8000 | 337 | 2400 |
| Site 2 | Siemens Verio 3T |  |  |  |  |  |
|  | 3D T1 weighted | 320 x 320 x 176 | 0.78 x 0.78 x 0.78 | 1900 | 3.4 | 900 |
|  | 3D T2 weighted | 512 x 512 x 160 | 0.488 x 0.488 x 1.0 | 3200 | 409 |  |
|  | FLAIR | 320 x 320 x 160 | 0.83 x 0.83 x 0.83 | 8000 | 337 | 2400 |
| Site 3 | Philips Achieva 3T |  |  |  |  |  |
|  | 3D T1 weighted | 320 x 320 x 200 | 0.8 x 0.8 x 0.8 | 6.2 | 3.0 |  |
|  | 3D T2 weighted | 320 x 320 x 200 | 0.8 x 0.8 x 0.8 | 2500 | 363 |  |
|  | FLAIR | 320 x 320 x 200 | 0.8 x 0.8 x 0.8 | 8000 | 337 | 2400 |
| Site 4 | Philips Achieva 3T |  |  |  |  |  |
|  | 3D T1 weighted | 320 x 320 x 112 | 0.8 x 0.8 x 1.6 | 6.5 | 3.2 |  |
|  | 3D T2 weighted | 320 x 320 x 112 | 0.8 x 0.8 x 1.6 | 2500 | 255 |  |
|  | FLAIR | 336 x 336 x 112 | 0.76 x 0.76 x 1.6 | 4800 | 330 | 1650 |
| Controls: Site1 | Philips 3T Achieva |  |  |  |  |  |
|  | 3D T1 weighted | 256 x 250 x 165 | 1.0 x 1.0 x 1.0 | 8.1 | 3.5 |  |
| Controls: OASIS | (Siemens TIM Trio |  |  |  |  |  |
|  | 3T) |  |  |  |  |  |
|  | 3D T1 weighted | 256 x 256 x 176 | 1.0 x 1.0 x 1.0 | 2400 | 3.16 |  |

3T = 3 Tesla; 3D=3-dimensional, TR = repetition time; TE = echo time; TI = inversion time; FLAIR = Fluid Attenuated Inversion Recovery; Site 1 control scans were acquired with parallel imaging, SENSE factor 1.5.

### Statistical Analyses

The analyses of neuropathological and morphological predictors of placebo response focused on MRI measures obtained at baseline, directly prior to extracranial venoplasty. Unless otherwise specified, all statistical analyses of morphometric and lesion data included age, gender, age by gender interaction, disease duration and site as covariates.

#### Placebo response

Change in MSQOL-54 Physical Health Composite was used as the measure of placebo response, as this measure showed a transient significant improvement in the trial (Traboulsee et al., 2018). Scores at baseline and the subsequent 6 assessment points were used to compute the area under the curve (AUC) measure for each participant using the Matlab function *trapz*. These values were then adjusted for the baseline score using linear regression, as higher baseline scores predicted less change (b = −0.91, SE = 0.32, t = −2.88, 95% CI = −1.53 - −0.28, p = 0.005), and standardized residuals were used to subdivide participants into placebo responders (scores > 0) and non-responders (scores ≤ 0).

#### Lesion analyses

These analyses aimed to determine whether placebo responders differed from non-responders in terms of lesion burden, type, and location.

Lesion segmentation was performed on FLAIR images by a trained radiologist blinded to treatment assignment using manual seed point and connected component analysis (McAusland, Tam, Wong, Riddehough, & Li, 2010). Lesion load (LL) was calculated as the sum of volumes of all FLAIR lesion masks (mm^3^) for a given patient. Lesion activity analysis was carried out by the same radiologist. Gadolinium enhanced lesions indicating blood brain barrier disruption were identified on T1 gadolinium scans. Newly enhancing lesions were counted as lesions that were enhanced on the current scan but were not enhancing in previous scans; new and newly enlarging lesions on the FLAIR scans were counted as lesions that were respectively absent or stable and smaller in volume in previous assessment points; unique newly active lesions were newly enhancing lesions and non-enhancing new or newly enlarging lesions on a current scan without double counting. Lesion load and its change over the trial’s duration was compared between placebo responders and non-responders with a linear mixed-effects model with our standard covariates using the lme4 R package (RRID:SCR_015654) (Bates, Mächler, Bolker, & Walker, 2015). Group differences in lesion counts at each assessment point were analyzed with Poisson regressions, as was change in lesion counts over time as a function of group.

The analysis of lesion location was performed in FSL (Functional MRI of the Brain Software Library, http://www.fmrib.ox.ac.uk/fsl, RRID:SCR_002823). Only the lesions identified on FLAIR scans were considered for this analysis, as there were too few participants with gadolinium enhanced lesions on the baseline scans (four non-responders; six responders, Table 4). FLAIR lesions masks were moved into the MNI-152 space, and lesion probability maps were created for the placebo responder and for the placebo non-responder groups by averaging the registered binary masks across patients in each group using the *fslmaths* utility. These voxel-wise maps representing the probability of each voxel being lesional were compared statistically between the two groups using the *Randomise* algorithm (Winkler, Ridgway, Webster, Smith, & Nichols, 2014), which uses non-parametric permutation inference to threshold a voxel-wise statistical map produced, in this case, by voxel-wise unpaired t-tests on the two groups.

#### MRI processing for brain morphometric and cortical thickness analyses

Patients’ normalized brain volumes (parenchymal volume normalized by intracranial volume) were computed on T1 MRIs using a segmentation-based approach (Wicks et al., 2015) and compared between placebo responders and non-responders using linear regression with our standard covariates. Percentage brain volume change was computed using an automated in-house method based on parenchymal edge displacement between scans (Smith et al., 2002); change over time was compared between responders and non-responders using a linear mixed-effects model (lme4 R package).

Prior to further morphometric analyses, white matter lesions on the patients’ scans were filled with intensities of neighboring white matter voxels using the *Lesion Filling* function in FSL (Battaglini, Jenkinson, & De Stefano, 2012). This reduces intensity contrast within lesion areas and can improve registration and segmentation of MS brains and resulting morphometric measurements (Valverde, Oliver, & Lladó, 2014). Native T1 MRIs were processed through the CIVET pipeline (version 2.1) (Zijdenbos, Forghani, & Evans, 2002) housed on the CBRAIN web-based image analysis platform (McGill Centre for Integrative Neuroscience, RRID:SCR_005513 (Sherif et al., 2014)). The pipeline included the following steps. 1) The native MRIs were registered to standard space using a 9-parameter nonlinear transformation (Collins, Neelin, Peters, & Evans, 1994). For the controls, the MNI-152 template was used, whereas for the MS patients, a high-resolution Alzheimer’s disease template provided by CIVET was used. Because of paraventricular brain atrophy in the MS patients, the AD template produced superior registration, and since our hypotheses pertained to the MS groups, we chose to use the template that was the most optimal for the patients. Simultaneous non-uniformity correction was performed using N3 algorithms (Sled, Zijdenbos, & Evans, 1998). 3) The registered corrected images were classified into white matter, gray matter, CSF, and background, using an artificial neural network classifier (Zijdenbos et al., 2002). 4) Binary volumes consisting of gray matter voxels and white matter voxels were then extracted from classified images, smoothed using an 8-mm FWHM smoothing kernel and used in the subsequent voxel-based morphometry analyses (see below). 5) Cortical surfaces were extracted from the classified images using the Constrained Laplacian Anatomic Segmentation Using Proximities (CLASP) surface extraction procedure (Kim et al., 2005)(Macdonald, Kabani, Avis, & Evans, 2000). This process generates a triangulated mesh at the interface of gray matter and white matter and expands the mesh outwards toward the pial surface. 6) Cortical thickness was measured in native space as the distance between corresponding vertices on the inner and outer surfaces of the mesh across 40,962 vertices in each hemisphere (Lerch & Evans, 2005). The cortical thickness maps were then blurred using a 20 mm surface based kernel (Chung et al., 2003). Quality control was performed using a combination of the CIVET QC tool and visual inspection of the outputs.

#### Voxel-based morphometry

Voxel-based morphometry was conducted on the 8-mm smoothed gray and white matter volumes using the PET and VBM module of Statistical Parametric Mapping (SPM 12, RRID:SCR_007037). The grey and white matter volumes were compared using voxel-wise two sample t-tests with our standard covariates plus total brain volume. Dimensional associations between volumes and the degree of placebo response, represented by the AUC change in MSQOL-54 scores adjusted for baseline scores, were also examined using multiple regression models with the same covariates. The resulting statistical maps were thresholded using family-wise error corrected threshold of p = 0.05.

#### Vertex-Wise Cortical thickness analysis

Native-space CT at 40,962 vertices was analyzed statistically using SurfStat (http://www.math.mcgill.ca/keith/surfstat/, RRID:SCR_007081), a Matlab toolbox for the statistical analysis of surface data with linear mixed effects models and random field theory to correct for multiple comparisons in determining vertex and cluster significance (Worsley, Andermann, Koulis, Macdonald, & Evans, 1999). The analyses were performed using Matlab17a. We first considered models predicting vertex-wise CT as a function of group membership (placebo responder vs. non-responder), adjusting for our standard covariates as well as total brain volume and lesion load at baseline:

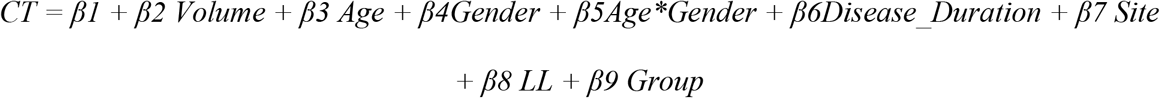

In light of the differential associations of lesion load with cortical thickness in placebo non-responders and responders (see Results), which may be viewed as violating the assumption of homogeneity of regression slopes (Miller & Chapman, 2001), the analysis was performed using models with and without lesion load as a covariate, as well as those substituting baseline EDSS as a measure of disease burden. As this did not change the results, we report the findings from the models with lesion load included as a covariate.

We additionally considered whether placebo response as a continuous variable was associated with CT. We tested this in a model predicting CT from the AUC measure of change in MSQOL-54, our standard covariates, total brain volume and lesion load at baseline and baseline MSQOL-54 scores:

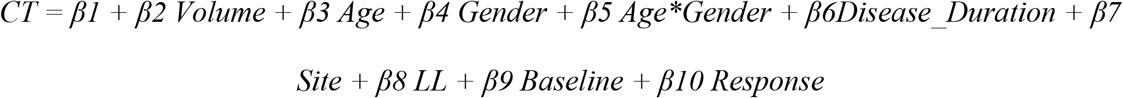

Given our finding of more regular CT graphs in placebo non-responders (see Results), which could arise from more coordinated tissue loss in anatomically connected regions owing to white matter lesions (He et al., 2009), we examined the associations between CT and lesion load in placebo responders versus non-responders. We first tested the significance of the interaction between lesion load and group in predicting CT.

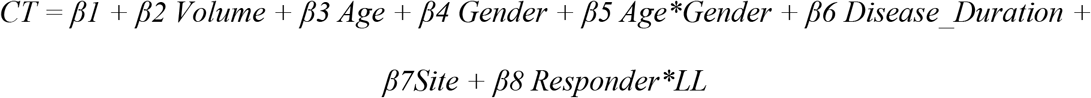

Once the significance of this interaction was established, we analyzed the associations between CT and lesion load separately in each group.

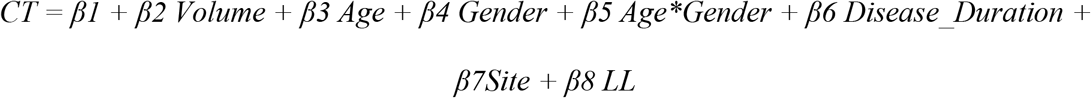

#### Graph theoretical cortical thickness analysis

Cortical surfaces were parcellated into 78 regions based on Automated Anatomical Labeling (AAL) (Tzourio-Mazoyer et al., 2002); subcortical labels were excluded. Mean CT values for each participant were extracted for each of the 78 regions. A linear regression was performed to adjust these CT values for total brain volume, age, gender and the interaction of age and gender for all participants, as well as disease duration, site and lesion load for the MS participants. Again, given differential associations of lesion load with cortical thickness in placebo non-responders and responders, the graph theoretical analyses were performed on the data both with and without lesion load covaried out, as well as on data adjusting for baseline EDSS in place of lesion load as a measure of disease burden. As this did not change the results, we report the results adjusted for lesion load. The resulting residuals were substituted for the raw cortical thickness values to construct inter-regional correlation matrices for placebo responders, non-responders and controls: Rij (i, j =1, 2 ... n, where n is the number of regions). These correlation matrices were then used to construct binarized networks and compute graph metrics at 30 linearly spaced sparsity thresholds ranging from 0.1 to 0.5. This thresholding approach normalizes each group-level graph to have the same number of edges. Using the Brain Connectivity Toolbox (RRID:SCR_004841) (Rubinov & Sporns, 2010) in Matlab2017a, we computed the following graph metrics: the clustering coefficient and its normalized version, the characteristic pathlength and its normalized version and the small-world index (Table 2).

**Table 2.**
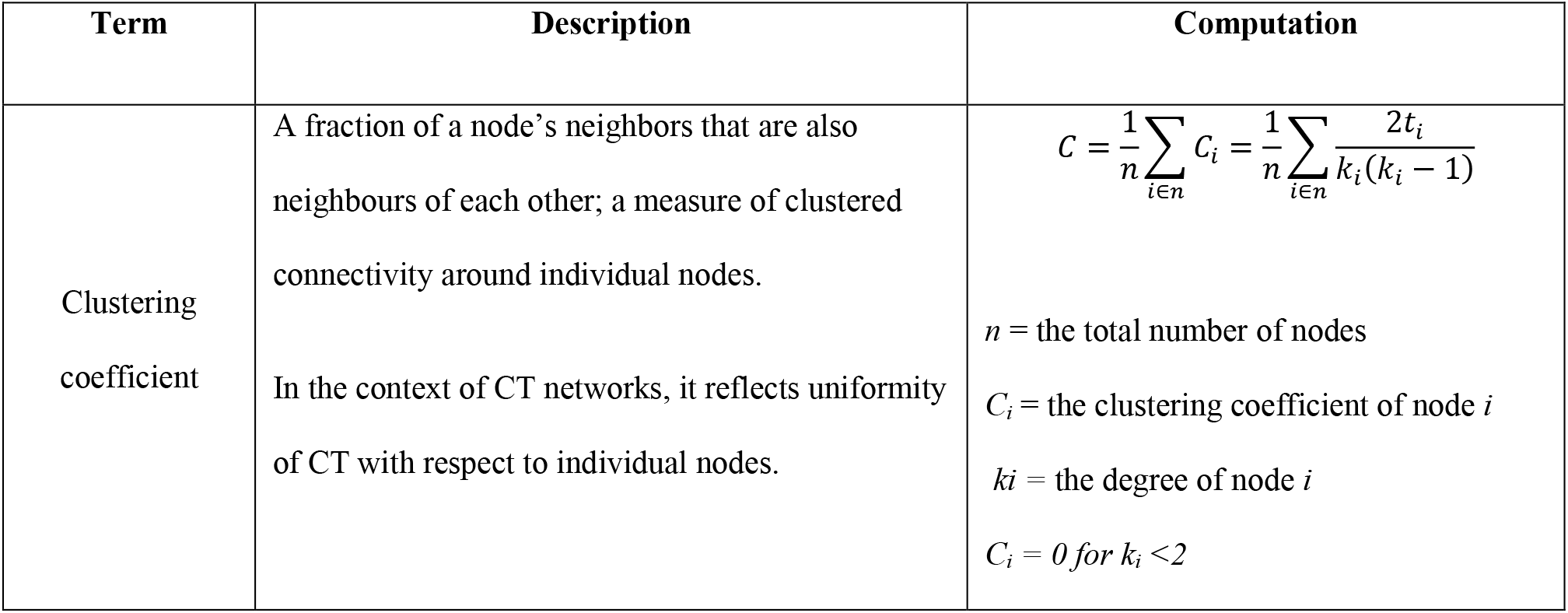
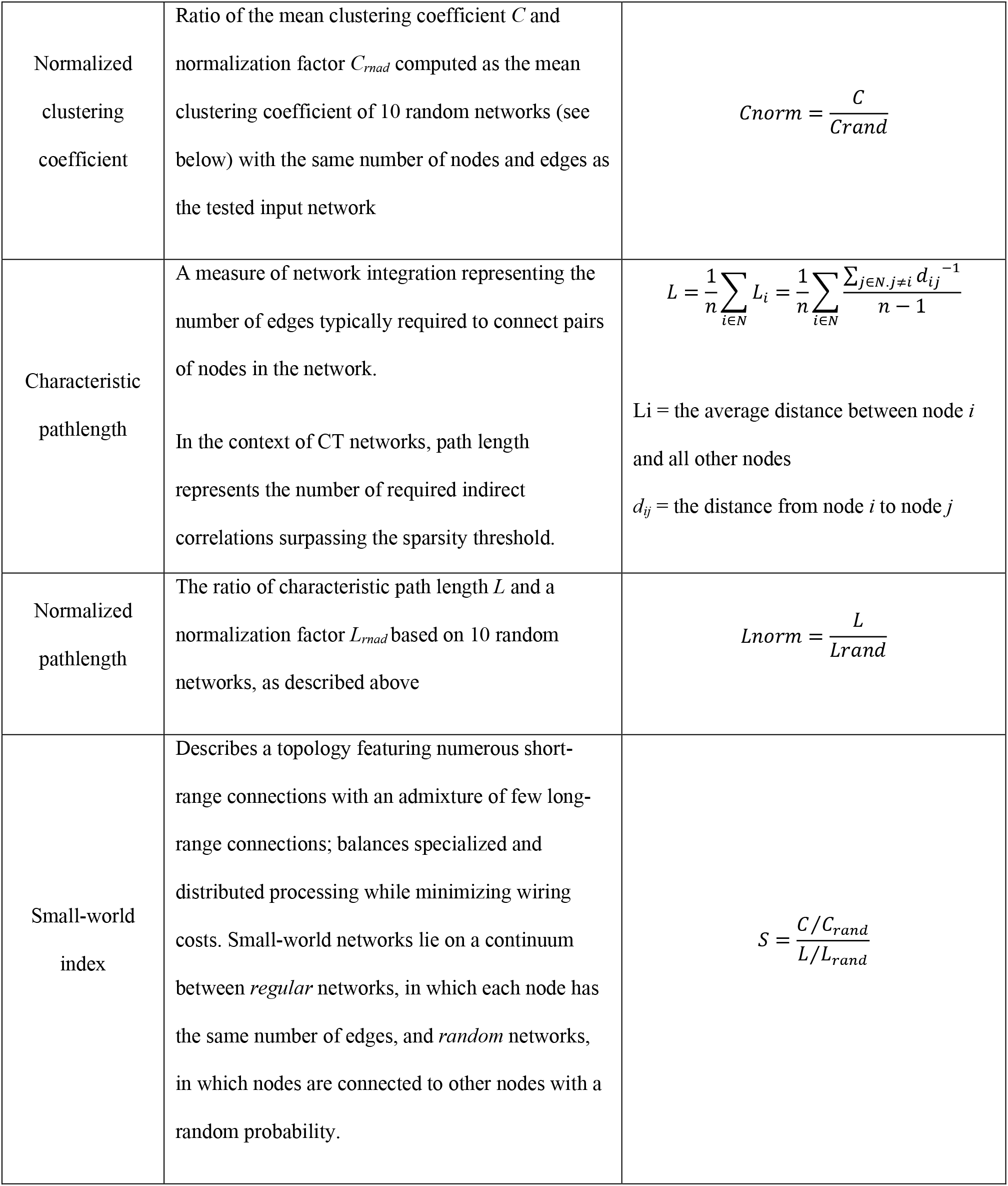
Graph theory metrics.

Leave-one-out cross-validation was performed to estimate the stability of the graph metrics, and error estimates from these cross validations were used to visualize group differences (Figure 2). Statistical significance of these differences was evaluated against null distributions of group differences on the metrics based on 1000 random permutations of participants in the input matrix. A Bonferroni correction for multiple comparisons was applied: considering three graph metrics (clustering coefficient, pathlength and small-world index) the α-level was determined to be 0.016. Because normalized and non-normalized variants of clustering coefficient and pathlength are redundant measures, these were not considered as separate comparisons.

**Figure 2.**
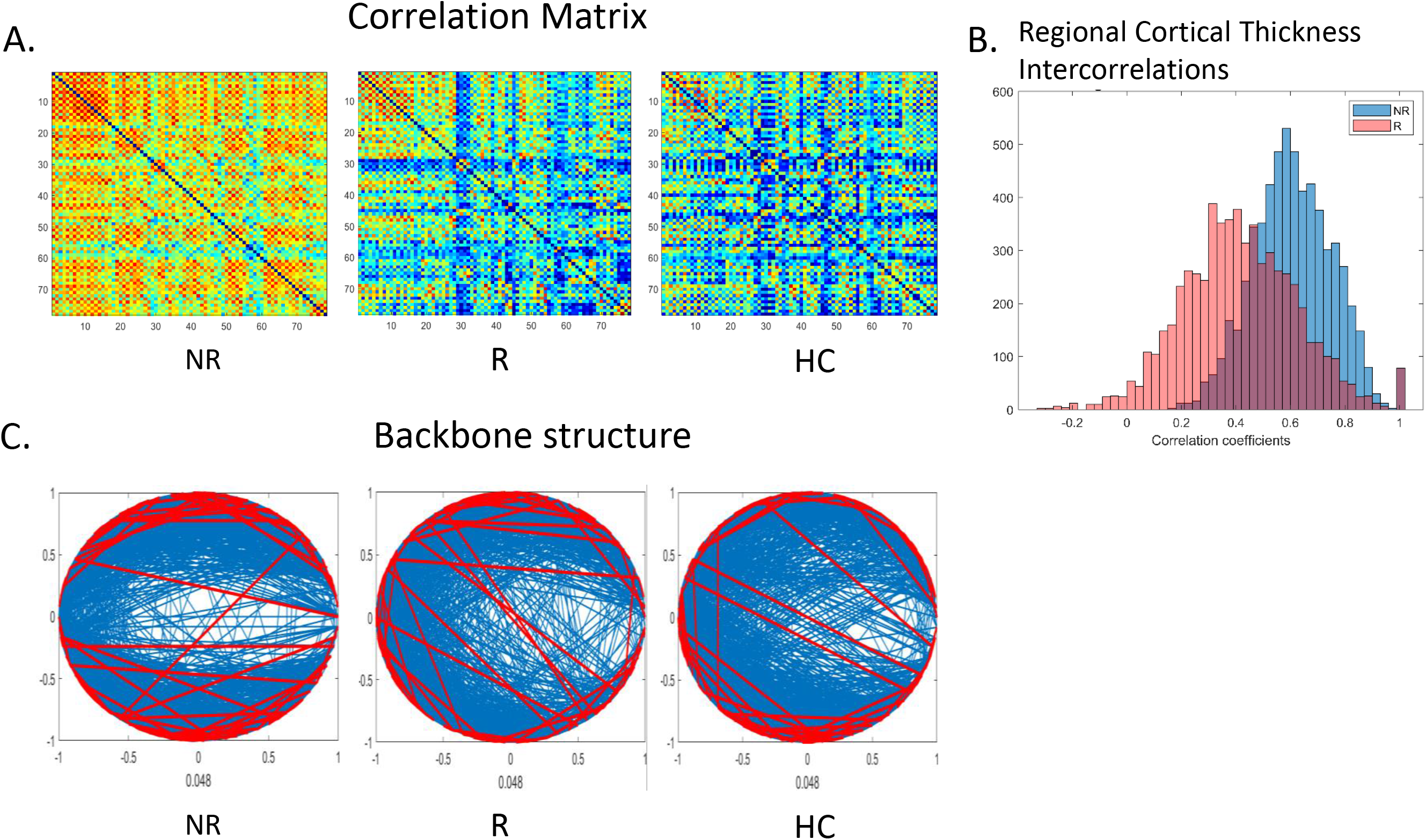
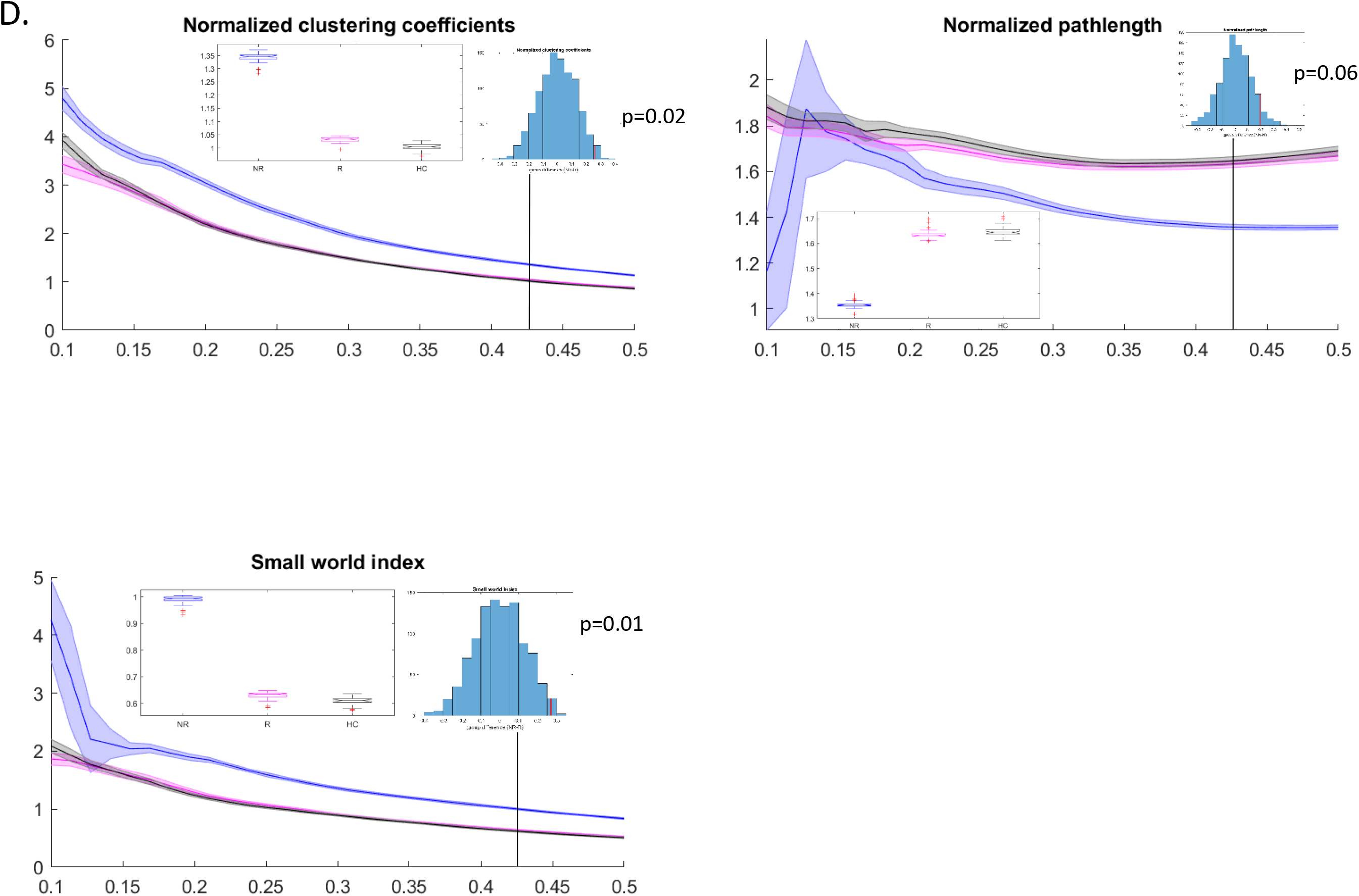
Cortical thickness covariance patterns in placebo responders and non-responders: graph theoretical analysis. (A) Correlation matrices of cortical thickness values across 78 cortical areas delineated using Automated Anatomical Labeling (AAL) in placebo non-responders (NR), placebo responders (R), and a group of healthy age-and gender-matched controls (HC). (B) A histogram depicting distributions of correlation coefficients in placebo responders and non-responders. (C) Backbone structure for correlation matrices in A at the sparsity threshold of 0.43. (D) Graph theoretical characteristics of these matrices across the range of sparsity thresholds from 0.1 to 0.5. Error ribbons represent standard deviation for parameter estimates from leave-one-out cross-validation. Box plots represent group comparisons at the sparsity threshold of 0.43 based on leave-one-out cross-validations; histograms represent the p-values based on permutation tests at the sparsity threshold of 0.43.

### Data Availability

Data and analysis code are available upon request.

## RESULTS

Placebo responders and non-responders did not differ significantly on any demographic or clinical characteristics (Table 3). The proportion of placebo responders vs. non-responders did not differ as a function of recruitment site. As expected, the two groups had markedly different subjective treatment response (Figure 1A).

**Table 3.** Demographic and clinical characteristics of placebo responders and non-responders.

|  | **Non-responder** | **Responder** | **p** |
| --- | --- | --- | --- |
|  | **(n = 45)** | **(n = 43)** |  |
| Venoplasty, n | 24 | 18 | 0.19 |
| Sham, n | 21 | 25 |  |
| Males, n | 18 | 13 | 0.24 |
| Females, n | 27 | 30 |  |
| Age, mean (SD) | 55.0 (7.4) | 53.0 (8.5) | 0.26 |
| Disease duration, mean (SD) | 18.8 (9.6) | 16.7 (8.1) | 0.29 |
| MS type (n) |  |  |  |
| RRMS | 25 | 30 |  |
| PPMS | 6 | 3 | 0.35 |
| SPMS | 14 | 10 |  |
| Baseline EDSS, median (range) | 4.0 (0-6.5) | 4.0 (0-6.5) | 0.09 |
| Baseline MSFC, mean (SD) | 7.2 (6.2) | 8.9 (4.9) | 0.17 |
| 25 foot walk, mean (SD) | 15.80 (12.15) | 11.87 (7.24) | 0.07 |
| 9 hole peg test, mean (SD) | 108.50 (49.73) | 94.41 (36.04) | 0.13 |
| PASAT, mean (SD) | 37.53 (14.70) | 38.63 (14.97) | 0.73 |
| Baseline MSQOL-54 PH | 55.54 (21.57) | 56.96 (18.79) | 0.74 |

RRMS = remitting relapsing MS; PPMS = primary progressive MS; SPMS = secondary progressive MS; EDSS = Expanded Disability Status Scale; MSFC = Multiple Sclerosis Functional Composite; PASAT = Paced Auditory Serial Addition Test; MSQOL-54 PH = Multiple Sclerosis Quality of Life – 54, Physical Health Composite. For MSFC, raw scores rather than z-scores are given.

Placebo responders and non-responders did not differ in terms of normalized brain volumes at baseline or percent brain volume change over the 48 weeks, although there was progressive atrophy in both groups (b = −0.31, SE = 0.09, t = −3.45, p = 0.0009). Voxel-based morphometry did not reveal any significant group differences in regional grey or white matter volume. Although there were no significant group differences in regional CT, there was a significant dimensional association between the magnitude of placebo response and CT of a left precuneus region (x = −3.16, y = −70.12, z = 37.10; Figure 3A).

**Figure 3:**
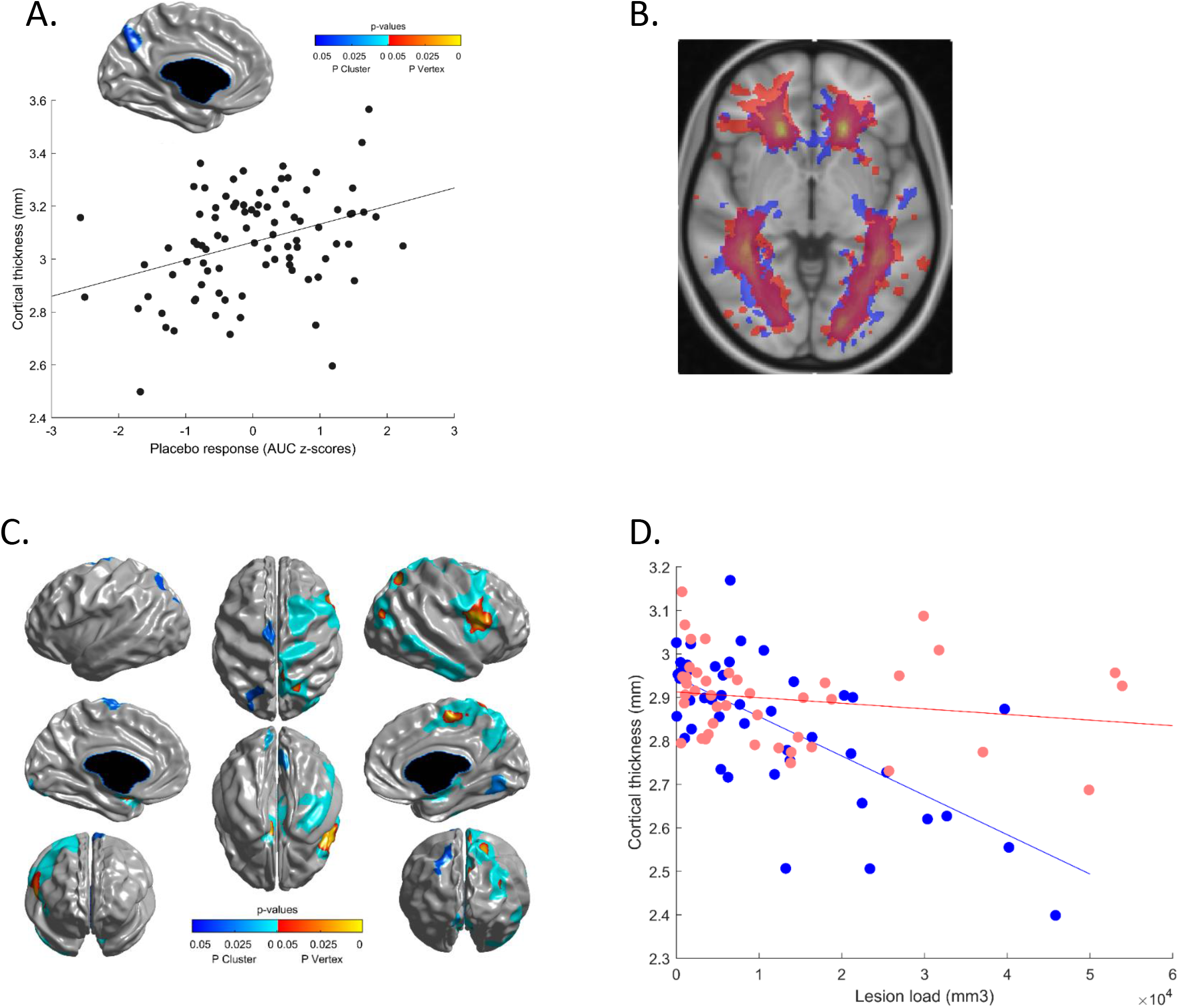
Cortical thickness, white matter lesions and placebo response. (A) Precuneus region whose thickness dimensionally predicts a stronger placebo response as depicted in the scatterplot. (B) Lesion probability maps in placebo responders (red) and non-responders (blue): no significant differences in lesion location between groups. (C) Cortical areas whose thickness was significantly associated with lesion load in the placebo non-responders. (D) Mean cortical thickness of the regions shown in relation to white matter lesion load (FLAIR) in the placebo responders (red) and non-responders (blue). A mask of the regions shown in A was used to extract mean CT values across these regions for all participants.

### Placebo Responders Have Higher Lesion Activity

Although most patients did not have gadolinium enhanced lesions, there were significant differences in lesion activity between the groups (Table 4). Placebo responders were more likely to display gadolinium enhanced lesions at baseline (b = 1.26, SE = 0.46, z = 2.73, p = 0.006) and to have newly enhancing lesions at 24 weeks (b = 1.26, SE = 0.42, z = 3.01, p = 0.003) and at 48 weeks (b = 1.77, SE = 0.51, z = 3.43, p = 0.0006). A similar pattern was evident for new lesions on FLAIR scans at both 24 (b = 1.11, SE = 0.33, z = 3.34, p = 0.0008) and 48 weeks (b = 1.57, SE = 0.40, z = 3.97, p = 0.00007) and for new active lesions that were not present on previous gadolinium T1 scans, both at 24 (b = 1.24, SE = 0.27, z = 4.52, p=0.000006) and 48 weeks (b = 1.54, SD = 0.36, z=4.25, p = 0.00002). The incidence of enlarging lesions was also higher in the placebo responders at 24 weeks (b = 2.61, SE = 0.75, z = 3.49, p = 0.0005), though not at 48 weeks (p = 0.14).

**Table 4.** Brain morphometric characteristics and lesions

| Measure (SD) | Non-responder (n = 45) | Responder (n = 43) | p |
| --- | --- | --- | --- |
| Normalized brain parenchymal fraction (baseline) | 0.75 (0.03) | 0.77 (0.03) | 0.39 |
| % Change in brain volume from baseline |  |  |  |
| 24 weeks | -0.40 (0.50) | -0.43 (0.38) | 0.78 |
| 48 weeks | -0.70 (0.60) | -0.61 (0.61) | 0.65 |
| WM lesion load (mm<sup>3</sup>) |  |  |  |
| Baseline | 12031.63 (12002.73) | 12269.53 (14538.12) | 0.78 |
| 24 weeks | 12681.97 (12185.17) | 12581.44 (14527.14) | 0.92 |
| 48 weeks | 12792.65 (11875.89) | 12907.21 (14698.09) | 0.87 |

| New T2 lesions (# of patients<br>with lesion count) | W24 | W48 | W24 | W48 |  |
| --- | --- | --- | --- | --- | --- |
| 0 | 36 | 36 | 35 | 33 |  |
| 1 | 4 | 5 | 3 | 4 |  |
| 2 | 3 | 0 | 1 | 3 |  |
| 3 | 0 | 2 | 0 | 1 |  |
| 5 | 1 | 0 | 0 | 0 |  |
| 6 | 0 | 0 | 1 | 0 | < 0.001** |
| 7 | 0 | 0 | 0 | 1 |  |
| 8 | 0 | 0 | 2 | 0 |  |
| 9 | 0 | 0 | 1 | 0 |  |
| 11 | 0 | 0 | 0 | 1 |  |

| Unique newly active lesions (#<br>of patients with lesion count) | BL | W24 | W48 | BL | W24 | W48 |  |
| --- | --- | --- | --- | --- | --- | --- | --- |
| 0 | 40 | 33 | 36 | 33 | 34 | 33 |  |
| 1 | 1 | 6 | 5 | 3 | 2 | 3 |  |
| 2 | 3 | 4 | 0 | 2 | 2 | 3 |  |
| 3 | 0 | 0 | 1 | 2 | 1 | 1 |  |
| 4 | 0 | 0 | 0 | 1 | 0 | 0 |  |
| 5 | 0 | 0 | 1 | 0 | 0 | 0 | < 0.0005*** |
| 6 | 0 | 1 | 0 | 0 | 1 | 1 |  |
| 7 | 0 | 0 | 0 | 1 | 0 | 0 |  |
| 8 | 0 | 0 | 0 | 0 | 1 | 1 |  |
| 11 | 0 | 0 | 0 | 0 | 0 | 1 |  |
| 15 | 0 | 0 | 0 | 0 | 1 | 0 |  |
| 22 | 0 | 0 | 0 | 0 | 1 | 0 |  |
| Newly enhancing lesions (# of patients with lesion count) | BL | W24 | W48 | BL | W24 | W48 |  |
| 0 | 40 | 36 | 38 | 33 | 36 | 34 |  |
| 1 | 1 | 6 | 4 | 3 | 4 | 5 |  |
| 2 | 3 | 2 | 1 | 2 | 0 | 0 |  |
| 3 | 0 | 0 | 0 | 2 | 1 | 2 |  |
| 4 | 0 | 0 | 0 | 1 | 0 | 0 |  |
| 5 | 0 | 0 | 0 | 0 | 0 | 2 | < 0.01* |
| 6 | 0 | 0 | 0 | 0 | 1 | 0 |  |
| 7 | 0 | 0 | 0 | 1 | 0 | 0 |  |
| Newly enlarging T2 lesions (# of patients with lesion count) | W24 | W48 | W24 | W48 |  |  |  |
| 0 | 42 | 42 | 38 | 38 |  |  |  |
| 1 | 2 | 0 | 2 | 4 |  |  | W24: 0.0005*** |
| 2 | 0 | 1 | 1 | 1 |  |  | W48: 0.14 |
| 7 | 0 | 0 | 1 | 0 |  |  |  |
| 13 | 0 | 0 | 1 | 0 |  |  |  |

BL = baseline; W24 = 24 weeks assessment point; W48 = 48 weeks assessment point; WM = white matter. P-values represent comparisons between placebo responders and non-responders for corresponding assessment points. The models include age, gender, age x gender interaction, disease duration and site as terms.

There was no significant change in lesion counts over the trial’s duration, and this did not differ as a function of group (p_s_ ≥ 0.2). There were no significant differences between placebo responders and non-responders in lesion load either at baseline or at 24-week and 48-week follow-up points (p_s_ ≥ 0.78), and there was no significant increase in lesion load over the 48 weeks (p = 0.11). There were also no significant differences in lesion locations (Figure 3B; supporting 3D movie).

### Placebo Non-Responders Have a More Homogenous and Small-World-Like CT Covariance Pattern

Group differences in the metrics presented below were consistent over the range of sparsity thresholds (10%-50% sparsity). As input matrices are not stable at low sparsity, which results in high error estimates, we chose a relatively high sparsity threshold of 43% yielding stable input matrices for presenting the results of random permutation tests. P-values for the entire range of sparsity thresholds are presented as supporting information (Table S1–S5); similar p-values were seen at most sparsity thresholds.

CT of placebo non-responders was more regionally homogeneous (range r: 0.1 to 0.95, median r: 0.66) relative to that of responders (range r_s_: −0.15 to 0.95, median r: 0.49) and controls (range r_s_: −0.22 to 0.92, median r: 0.41, Figure 2A, B). Placebo responders did not differ significantly from controls on the computed metrics (p_s_ ≥ 0.29). Uncorrected results showed that non-responders had a more segregated network topology with higher mean and normalized clustering coefficients (non-responders vs. responders: p = 0.02; non-responders vs. controls: p = 0.01). Non-responders also had marginally shorter pathlengths relative to responders (p = 0.06) and controls (p_s_ < 0.04). This resulted in stronger small-world attributes for non-responders relative to responders and controls (p_s_ = 0.01), indicating a shift towards more regular and less random graphs. The group differences in small world index and the difference between non-responders and controls in clustering coefficient survived the correction for multiple comparisons.

### Lesion Load Inversely Predicts CT Only in Placebo Non-Responders

Although lesion load and location did not differ significantly between responders and non-responders, there was a significant difference between the groups in the association between CT and lesion load. While there was no relationship between lesion load and CT in responders, in non-responders, greater lesion load was associated with cortical thinning in 9 clusters (Figure 3C, D, Table 5). The clusters covered a substantial portion of the right hemisphere including parts of the primary motor and sensory cortices as well as the premotor cortex and somatosensory, visual and auditory association areas. The major clusters included: the primary motor cortex, comprising both the paracentral lobule and the precentral gyrus and extending into the premotor cortex (middle frontal gyrus, Brodmann area BA 6) and the insula; primary somatosensory cortex (postcentral gyrus, BA 3) extending to the superior parietal lobule and the precuneus; superior occipital gyrus (BA 19) extending to middle temporal gyrus (BA 39); middle temporal gyrus (BA 21) extending to inferior temporal gyrus (BA 20). There were also smaller primary motor and superior occipital clusters in the left hemisphere.

**Table 5:** Regions where cortical thickness was predicted by lesion load in placebo non-responders.

| Cluster | # vertices | Peak | x | y | z | t | p |
| --- | --- | --- | --- | --- | --- | --- | --- |
| Right superior and medial frontoparietal<br>$p < 0.00005$ | 4080 | Medial frontal gyrus (R) | 4.30 | -24.62 | 57.64 | 5.22 | 0.008 |
|  |  | Superior parietal lobule (R) | 28.69 | -57.71 | 52.84 | 5.03 | 0.013 |
|  |  | Precuneus (R) | 13.35 | -66.54 | 32.83 | 4.96 | 0.015 |
|  |  | Middle occipital gyrus (R) | 44.45 | -77.33 | 22.46 | 4.83 | 0.021 |
|  |  | Postcentral gyrus (R) | 11.82 | -59.08 | 69.79 | 4.75 | 0.027 |
|  |  | Paracentral lobule (R) | 13.69 | -41.22 | 58.15 | 4.56 | 0.043 |
| Right precentral gyrus / insula<br>$p < 0.00005$ | 3026 | Precentral gyrus (R) | 52.78 | -6.92 | 10.30 | 5.32 | 0.006 |
|  |  | Insula (R) | 39.91 | -0.08 | 16.26 | 5.21 | 0.008 |
| Left parahippocampal gyrus<br>$p < 0.00005$ | 383 | Parahippocampal gyrus (L) | -12.13 | -9.54 | -12.02 | 5.41 | 0.005 |
| Right inferior temporal gyrus<br>$p = 0.0002$ | 1220 | Inferior temporal gyrus (R) | 59.56 | -7.57 | -28.66 | 4.51 | 0.05 |
| Right parahippocampal gyrus<br>$p = 0.0005$ | 335 | Parahippocampal gyrus (R) | 15.12 | -12.79 | -11.08 | 4.82 | 0.02 |
| Left lingual gyrus<br>$p = 0.01$ | 246 | - | -16.26 | -93.19 | -12.51 | 3.14 | - |
| Right lingual gyrus<br>p = 0.02 | 311 | - | 10.08 | -72.09 | -4.17 | 4.03 | - |
| Left precentral gyrus<br>p = 0.03 | 385 | - | -14.29 | -19.43 | 70.91 | 4.12 | - |
| Left superior parietal lobule<br>p = 0.03 | 329 | - | -31.33 | -70.83 | 47.50 | 3.64 | - |

## DISCUSSION

We examined neural correlates of placebo responses to an ineffective invasive procedure, which nonetheless inspired hope in patients, driving many to pursue it despite the potential risks. Our findings revealed significant structural brain differences between the MS patients who experienced these real-world placebo responses and those who did not. These findings advance our understanding of both the neurobiology of placebo responses and of how the neuropathological changes in MS impact the propensity to experience them.

While placebo responders and non-responders did not differ significantly in terms of clinical characteristics, lesion load, lesion location, brain volume or regional cortical thickness, the groups differed significantly in terms of their CT covariance patterns. Relative to placebo responders, non-responders had more uniform CT across different brain regions with a more segregated and clustered topology. Coupled with marginally shorter pathlengths, this resulted in stronger small-world attributes, indicating a shift towards more regular and less random graphs. While a more regular network may be associated with smaller wiring costs, potentially imposed by axonal loss in MS, this type of network may be less capable of distributed processing and functional integration. The absence of differences between responders and controls, suggests that the more segregated and regular topology observed in non-responders is anomalous.

Previous studies have suggested that CT covariance networks in MS are characterized by increased segregation with an enhancement of local properties, as well as a shift towards more regular networks in advanced disease (Fleischer et al., 2019)(Muthuraman et al., 2016)(He et al., 2009)(Tewarie et al., 2014). Convergent findings have emerged from studies of DTI-based structural networks (Fleischer et al., 2017) (Shu et al., 2011) and MEG-based functional connectivity in MS patients (Tewarie et al., 2014). Together, the evidence points to a pathological shift towards more segregated and regular networks in MS, and our findings suggest that the MS patients who experience these shifts may also lose their capacity to experience placebo effects. It follows that placebo responses may require a cortical network topology that favours distributed processing and functional integration.

One interpretation of our findings is that in placebo non-responders, the disease process may have resulted in more synchronized cortical tissue loss across different brain regions leading to increasingly correlated cortical thickness values. This interpretation is supported by the associations we found between lesion load and regional CT in placebo non-responders only. Although white matter demyelination is thought to drive neuronal degeneration in MS, there is evidence that the two can occur independently, and laminar contributions to cortical neuronal loss (and hence thinning) may differ depending on whether it is related to versus independent from white matter demyelination (Trapp et al., 2018). In the non-responders, lesion load significantly predicted cortical thinning in a substantial portion of the right hemisphere, including primary sensory and motor areas and somatosensory, visual and auditory association areas. Together with precuneus and insula, association areas have been identified as hubs by graph theoretical studies of structural connectivity (van den Heuvel & Sporns, 2013). Insults to these hubs and their connections are likely to result in changes in network organization with major implications for functional integration of neural activity (Gratton, Nomura, Pérez, & D’Esposito, 2012) (Crossley et al., 2014). Synchronous loss of neurons in these regions and of projections between them could impair associative processes enabling placebo responses, such as integrating interoceptive appraisals with expectancy of therapeutic benefit. Precuneus and insula, which are key structures for self-referential thinking and interoceptive awareness, may play a central role in such expectancy-informed appraisals. Loss of projections between these regions and shared deep nuclei could also result in an organization increasingly dependent on local connections.

Although the causes of the differential associations between CT and lesion load in placebo responders versus non-responders remain undetermined, lesion characteristics may play a role. Cortical grey matter loss in MS may arise from a combination of primary pathological processes and secondary effects of white matter damage. Regarding the latter, chronic inactive lesions are more likely to be associated with axonal degeneration (Mahad et al., 2015). Although both placebo responders and non-responders had relatively advanced disease (10-20 years, median EDSS of 4.0), at which point most lesions are inactive (Frischer et al., 2015), and gadolinium enhancing lesions were observed in a minority of patients, responders had a significantly higher incidence of active lesions. The same subset of patients also had more lesions that became enlarged at 24 or 48 weeks relative to baseline, suggesting either expanding inflammatory activity or slowly expanding “smoldering” lesions (Frischer et al., 2015). Based on this, and given equivalent lesion load in the two groups, it is plausible that placebo non-responders had a higher proportion of inactive lesions, more likely to be chronic and to reflect axonal loss potentially driving synchronized loss of cortical tissue. In addition, the absence of active inflammation in non-enhanced lesions may have caused their volume to be reduced compared to that of active lesions, resulting in an apparently smaller lesion load. Lesion location, on the other hand, did not drive differential associations between lesion load and CT, as lesion maps did not differ between responders and non-responders.

Whether the network characteristics we observed to be associated with the absence of placebo response would generalize to other types of placebo responses or other patient groups remains to be determined. This could be tested by applying graph theoretical analysis in future studies of the neural mechanisms of placebo responses or to existing published datasets. Thus far, graph theory has seldom been used to study neural correlates of placebo responses. We are aware of only one study using graph theory metrics of DTI-based structural networks to predict placebo response in migraine patients (Liu et al., 2017). In that study, increased global and local efficiency at baseline inversely predicted placebo analgesic response to sham acupuncture, which is broadly consistent with our findings.

Our study had several limitations. First, to maximize sample size, sham and venoplasty participants were combined under the reasonable assumption that both interventions were effectively sham, considering that venoplasty for MS had a questionable scientific rationale and was found ineffective in two independent trials. Indeed, the responder group included non-significantly more sham participants (Table 3). Moreover, supplementary analyses performed separately in sham and venoplasty groups yielded similar trends (Supporting information, Tables S4–S5), highlighting the robustness of the differences in CT networks between responders and non-responders. We consider the findings from the full sample more likely to be reliable.

Second, we excluded some patients due to poor MRI quality. Third, the control scans were acquired on a different scanner than the patient scans, which could have introduced a bias. However, the primary comparisons were between the two patient groups, and the control group was used to provide a benchmark for interpreting the graphs derived from the placebo responder and non-responder groups. Fourth, responders and non-responders were identified based on subjective self-report of health-related quality of life. Hence, our findings may not generalize to placebo responses manifesting as more objective clinical improvement, which we did not observe in the trial (Traboulsee et al., 2018). However, even subjective placebo responses can be quite compelling for the patients: in the case of venoplasty, they may have contributed to fueling the efforts of patient advocacy groups to legitimize the procedure in the face of skepticism from the scientific community (Paylor et al., 2014). Fifth, in a trial with a 48-week follow-up, placebo response is necessarily confounded with the natural course of the disease including relapses, remissions and regression to the mean. We consider substantial effects of such confounds unlikely because 1) we adjusted our measure of placebo response for baseline scores to minimize the impact of regression to the mean; and 2) the incidence of active lesions did not change in either group over the course of the trial, making it unlikely that placebo response was driven by remissions. The latter is further supported by the absence placebo response on more objective clinician-rated measures, such as the EDSS. Finally, our graph theoretical analysis was based on cortical thickness covariance patterns, which do not represent either true structural connectivity or a direct measure of functional connectivity. Rather, they are thought of as an indirect reflection of functional connectivity between brain regions. Resting state fMRI data could have provided valuable direct information regarding functional connectivity differences between placebo responders and non-responders. Unfortunately, no resting state sequences were collected as part of this trial. Although DTI sequences were available, DTI as measure of structural connectivity is problematic in advanced MS, as white matter lesions present a challenge for tract identification.

In conclusion, our findings demonstrate that the absence of placebo response in MS is associated with 1) a more regular and segregated CT topology, 2) cortical tissue loss related to white matter pathology, and 3) lower lesion activity. Considering that placebo response is a constituent of active therapeutic response, these morphometric characteristics may by extension predict responses to active therapies. Finally, our findings highlight graph theory as a promising tool for future studies of the neurobiology of placebo responses.

---

**Table S1.** Permutation test p-values for placebo responders versus non-responders at different sparsity thresholds.

| Sparsity threshold | Mean clustering coefficient | Normalized clustering coefficient | Characteristic pathlength | Normalized pathlength | Small world index |
| --- | --- | --- | --- | --- | --- |
| 0.10 | **0.00\*** | **0.02\*** | 0.06† | 0.06† | **0.01\*** |
| 0.11 | **0.01\*** | **0.03\*** | **0.021\*** | **0.02\*** | **0.00\*** |
| 0.13 | **0.02\*** | **0.02\*** | 0.51 | 0.50 | 0.21 |
| 0.14 | **0.04\*** | **0.03\*** | 0.54 | 0.53 | 0.20 |
| 0.16 | **0.04\*** | 0.06† | 0.52 | 0.52 | 0.14 |
| 0.17 | 0.07† | **0.04\*** | 0.46 | 0.46 | 0.07† |
| 0.18 | 0.07† | 0.06† | 0.39 | 0.39 | **0.04\*** |
| 0.20 | 0.13 | 0.19 | 0.40 | 0.40 | 0.09† |
| 0.21 | **0.05\*** | 0.07† | 0.40 | 0.40 | **0.03\*** |
| 0.22 | **0.03\*** | **0.04\*** | 0.31 | 0.31 | **0.01\*** |
| 0.24 | **0.03\*** | **0.03\*** | 0.25 | 0.24 | **0.01\*** |
| 0.25 | **0.04\*** | **0.03\*** | 0.22 | 0.22 | **0.01\*** |
| 0.27 | **0.04\*** | **0.04\*** | 0.21 | 0.21 | **0.01\*** |
| 0.28 | **0.02\*** | **0.02\*** | 0.19 | 0.19 | **<0.001\*** |
| 0.29 | **0.02\*** | **0.02\*** | 0.16 | 0.16 | **<0.001\*** |
| 0.31 | **0.02\*** | **0.03\*** | 0.15 | 0.15 | **<0.001\*** |
| 0.32 | **0.02\*** | **0.02\*** | 0.12 | 0.12 | **<0.001\*** |
| 0.33 | **0.02\*** | **0.02\*** | 0.11 | 0.11 | **0.01\*** |
| 0.35 | **0.03\*** | **0.02\*** | 0.10 | 0.10 | **0.01\*** |
| 0.36 | **0.02\*** | **0.02\*** | 0.09† | 0.09† | **0.01\*** |
| 0.38 | **0.02\*** | **0.02\*** | 0.08† | 0.08† | **0.01\*** |
| 0.39 | **0.02\*** | **0.02\*** | 0.07† | 0.07† | **0.01\*** |
| 0.40 | **0.02\*** | **0.02\*** | 0.07† | 0.07† | **0.01\*** |
| 0.42 | **0.02\*** | **0.02\*** | 0.06† | 0.06† | **0.01\*** |
| 0.43 | **0.02\*** | **0.02\*** | 0.06† | 0.06† | **0.01\*** |
| 0.44 | **0.02\*** | **0.02\*** | 0.05† | 0.05† | **0.01\*** |
| 0.46 | **0.02\*** | **0.03\*** | 0.05† | 0.05† | **0.03\*** |
| 0.47 | **0.05\*** | **0.04\*** | 0.05† | 0.05† | **0.03\*** |
| 0.49 | **0.05\*** | **0.05\*** | 0.05† | 0.05† | **0.04\*** |
| 0.50 | **0.03\*** | **0.03\*** | 0.05† | 0.06† | **0.03\*** |

\* $p \leq 0.05$ ; † $p \leq 0.1$

**Table S2.** Permutation test p-values for controls versus placebo non-responders at different sparsity thresholds.

| Sparsity threshold | Mean clustering coefficient | Normalized clustering coefficient | Characteristic pathlength | Normalized pathlength | Small world index |
| --- | --- | --- | --- | --- | --- |
| 0.10 | 0.09† | **0.05\*** | 0.06† | 0.08† | **0.03\*** |
| 0.11 | 0.06† | **0.05\*** | 0.49 | 0.06† | **0.01\*** |
| 0.13 | 0.07† | 0.23 | 0.50 | 0.49 | 0.41 |
| 0.14 | 0.08† | 0.14 | 0.49 | 0.50 | 0.34 |
| 0.16 | 0.15 | 0.15 | 0.50 | 0.49 | 0.28 |
| 0.17 | 0.11 | 0.12 | 0.40 | 0.50 | 0.23 |
| 0.18 | 0.11 | 0.09† | 0.42 | 0.40 | 0.10 |
| 0.20 | 0.11 | 0.14 | 0.40 | 0.42 | 0.09† |
| 0.21 | **0.04\*** | 0.07† | 0.35 | 0.40 | **0.04\*** |
| 0.22 | **0.05\*** | **0.03\*** | 0.32 | 0.35 | **0.02\*** |
| 0.24 | **0.05\*** | 0.05† | 0.31 | 0.32 | **0.02\*** |
| 0.25 | **0.05\*** | 0.05† | 0.25 | 0.31 | **0.02\*** |
| 0.27 | **0.04\*** | **0.03\*** | 0.20 | 0.26 | **0.01\*** |
| 0.28 | **0.04\*** | **0.05\*** | 0.18 | 0.20 | **0.02\*** |
| 0.29 | 0.06† | 0.08† | 0.17 | 0.18 | **0.02\*** |
| 0.31 | 0.06† | 0.05† | 0.14 | 0.17 | **0.02\*** |
| 0.32 | **0.04\*** | **0.05\*** | 0.11 | 0.14 | **0.02\*** |
| 0.33 | **0.05\*** | **0.04\*** | 0.11 | 0.11 | **0.02\*** |
| 0.35 | 0.06† | 0.06† | 0.08† | 0.11 | **0.03\*** |
| 0.36 | **0.04\*** | **0.05\*** | 0.06† | 0.08† | **0.02\*** |
| 0.38 | **0.04\*** | **0.03\*** | **0.05\*** | 0.06† | **0.02\*** |
| 0.39 | **0.02\*** | **0.03\*** | **0.05\*** | 0.05† | **0.02\*** |
| 0.40 | **0.02\*** | **0.02\*** | **0.05\*** | **0.05\*** | **0.02\*** |
| 0.42 | **0.01\*** | **0.01\*** | **0.04\*** | **0.05\*** | **0.01\*** |
| 0.43 | **0.01\*** | **0.01\*** | **0.03\*** | **0.04\*** | **0.01\*** |
| 0.44 | **0.01\*** | **0.01\*** | **0.03\*** | **0.03\*** | **0.01\*** |
| 0.46 | **0.01\*** | **0.01\*** | **0.03\*** | **0.03\*** | **0.01\*** |
| 0.47 | **0.02\*** | **0.02\*** | **0.03\*** | **0.03\*** | **0.02\*** |
| 0.49 | **0.02\*** | **0.02\*** | **0.03\*** | **0.03\*** | **0.02\*** |
| 0.50 | **0.02\*** | **0.02\*** | 0.07† | **0.03\*** | **0.02\*** |

\* $p \leq 0.05$ ; † $p \leq 0.1$

**Table S3.** Permutation test p-values for controls versus placebo responders at different sparsity thresholds

| Sparsity threshold | Mean clustering coefficient | Normalized clustering coefficient | Characteristic pathlength | Normalized pathlength | Small world index |
| --- | --- | --- | --- | --- | --- |
| 0.10 | 0.83 | 0.50 | 0.26 | 0.27 | 0.35 |
| 0.11 | 0.66 | 0.54 | 0.48 | 0.49 | 0.51 |
| 0.13 | 0.67 | 0.63 | 0.41 | 0.43 | 0.55 |
| 0.14 | 0.57 | 0.59 | 0.44 | 0.42 | 0.53 |
| 0.16 | 0.71 | 0.73 | 0.43 | 0.43 | 0.65 |
| 0.17 | 0.59 | 0.69 | 0.58 | 0.57 | 0.71 |
| 0.18 | 0.60 | 0.62 | 0.38 | 0.39 | 0.52 |
| 0.20 | 0.39 | 0.43 | 0.38 | 0.37 | 0.36 |
| 0.21 | 0.39 | 0.42 | 0.38 | 0.38 | 0.37 |
| 0.22 | 0.51 | 0.53 | 0.42 | 0.42 | 0.48 |
| 0.24 | 0.45 | 0.54 | 0.47 | 0.47 | 0.51 |
| 0.25 | 0.41 | 0.38 | 0.52 | 0.53 | 0.42 |
| 0.27 | 0.40 | 0.40 | 0.45 | 0.45 | 0.40 |
| 0.28 | 0.59 | 0.56 | 0.43 | 0.43 | 0.50 |
| 0.29 | 0.65 | 0.61 | 0.43 | 0.43 | 0.55 |
| 0.31 | 0.62 | 0.55 | 0.42 | 0.42 | 0.49 |
| 0.32 | 0.55 | 0.51 | 0.45 | 0.45 | 0.48 |
| 0.33 | 0.43 | 0.48 | 0.43 | 0.43 | 0.45 |
| 0.35 | 0.47 | 0.45 | 0.43 | 0.43 | 0.44 |
| 0.36 | 0.46 | 0.46 | 0.40 | 0.40 | 0.42 |
| 0.38 | 0.42 | 0.42 | 0.39 | 0.39 | 0.40 |
| 0.39 | 0.34 | 0.33 | 0.41 | 0.41 | 0.36 |
| 0.40 | 0.33 | 0.33 | 0.40 | 0.40 | 0.35 |
| 0.42 | 0.30 | 0.31 | 0.40 | 0.40 | 0.34 |
| 0.43 | 0.30 | 0.33 | 0.39 | 0.39 | 0.35 |
| 0.44 | 0.30 | 0.33 | 0.38 | 0.38 | 0.34 |
| 0.46 | 0.29 | 0.29 | 0.37 | 0.37 | 0.32 |
| 0.47 | 0.28 | 0.28 | 0.35 | 0.35 | 0.31 |
| 0.49 | 0.27 | 0.26 | 0.35 | 0.35 | 0.30 |
| 0.50 | 0.31 | 0.32 | 0.34 | 0.34 | 0.33 |

\* $p \leq 0.05$ ; † $p \leq 0.1$

**Table S4.** Permutation test p-values for placebo responders versus non-responders including only sham-treated participants at different sparsity thresholds.

| Sparsity threshold | Mean clustering coefficient | Normalized clustering coefficient | Characteristic pathlength | Normalized pathlength | Small world index |
| --- | --- | --- | --- | --- | --- |
| 0.10 | **0.01\*** | **0.02\*** | 0.82 | 0.83 | 0.33 |
| 0.11 | **0.01\*** | **0.01\*** | 0.84 | 0.85 | 0.27 |
| 0.13 | **<0.00\*** | **<0.00\*** | 0.83 | 0.83 | 0.12 |
| 0.14 | **<0.00\*** | **0.01\*** | 0.86 | 0.86 | 0.16 |
| 0.16 | **0.01\*** | **0.02\*** | 0.93 | 0.93 | 0.35 |
| 0.17 | **<0.001\*** | **<0.001\*** | 0.91 | 0.91 | 0.16 |
| 0.18 | **<0.001\*** | **<0.001\*** | 0.79 | 0.79 | **0.05\*** |
| 0.20 | **<0.001\*** | **<0.001\*** | 0.73 | 0.73 | **0.03\*** |
| 0.21 | **<0.001\*** | **<0.001\*** | 0.74 | 0.74 | **0.03\*** |
| 0.22 | **0.03\*** | **0.04\*** | 0.68 | 0.68 | 0.10 |
| 0.24 | **0.04\*** | **0.04\*** | 0.65 | 0.65 | 0.08† |
| 0.25 | **0.04\*** | **0.04\*** | 0.59 | 0.58 | 0.07† |
| 0.27 | 0.06† | **0.10\*** | 0.59 | 0.59 | 0.11 |
| 0.28 | 0.06† | 0.08† | 0.55 | 0.55 | 0.10 |
| 0.29 | 0.06† | **0.05\*** | 0.45 | 0.45 | 0.06† |
| 0.31 | 0.08† | 0.07† | 0.43 | 0.43 | 0.08† |
| 0.32 | 0.07† | 0.07† | 0.39 | 0.39 | 0.08† |
| 0.33 | **0.05\*** | 0.06† | 0.42 | 0.42 | 0.07† |
| 0.35 | **0.05\*** | **0.05\*** | 0.36 | 0.36 | 0.07† |
| 0.36 | **0.03\*** | **0.03\*** | 0.27 | 0.27 | **0.04\*** |
| 0.38 | **0.04\*** | **0.04\*** | 0.26 | 0.26 | **0.04\*** |
| 0.39 | **0.04\*** | **0.04\*** | 0.25 | 0.25 | 0.06† |
| 0.40 | **0.04\*** | **0.04\*** | 0.25 | 0.25 | 0.06† |
| 0.42 | **0.05\*** | **0.05\*** | 0.23 | 0.23 | 0.07† |
| 0.43 | 0.06† | 0.06† | 0.24 | 0.24 | 0.08† |
| 0.44 | 0.06† | 0.07† | 0.24 | 0.24 | 0.09† |
| 0.46 | 0.07† | 0.07† | 0.23 | 0.23 | 0.09† |
| 0.47 | 0.08† | 0.09† | 0.22 | 0.22 | 0.11 |
| 0.49 | 0.11 | 0.11 | 0.22 | 0.22 | 0.13 |
| 0.50 | 0.13 | 0.12 | 0.22 | 0.22 | 0.15 |

\*p ≤ 0.05; †p ≤ 0.1; sham responders: n = 25; sham non-responders: n = 21

**Table S5:**
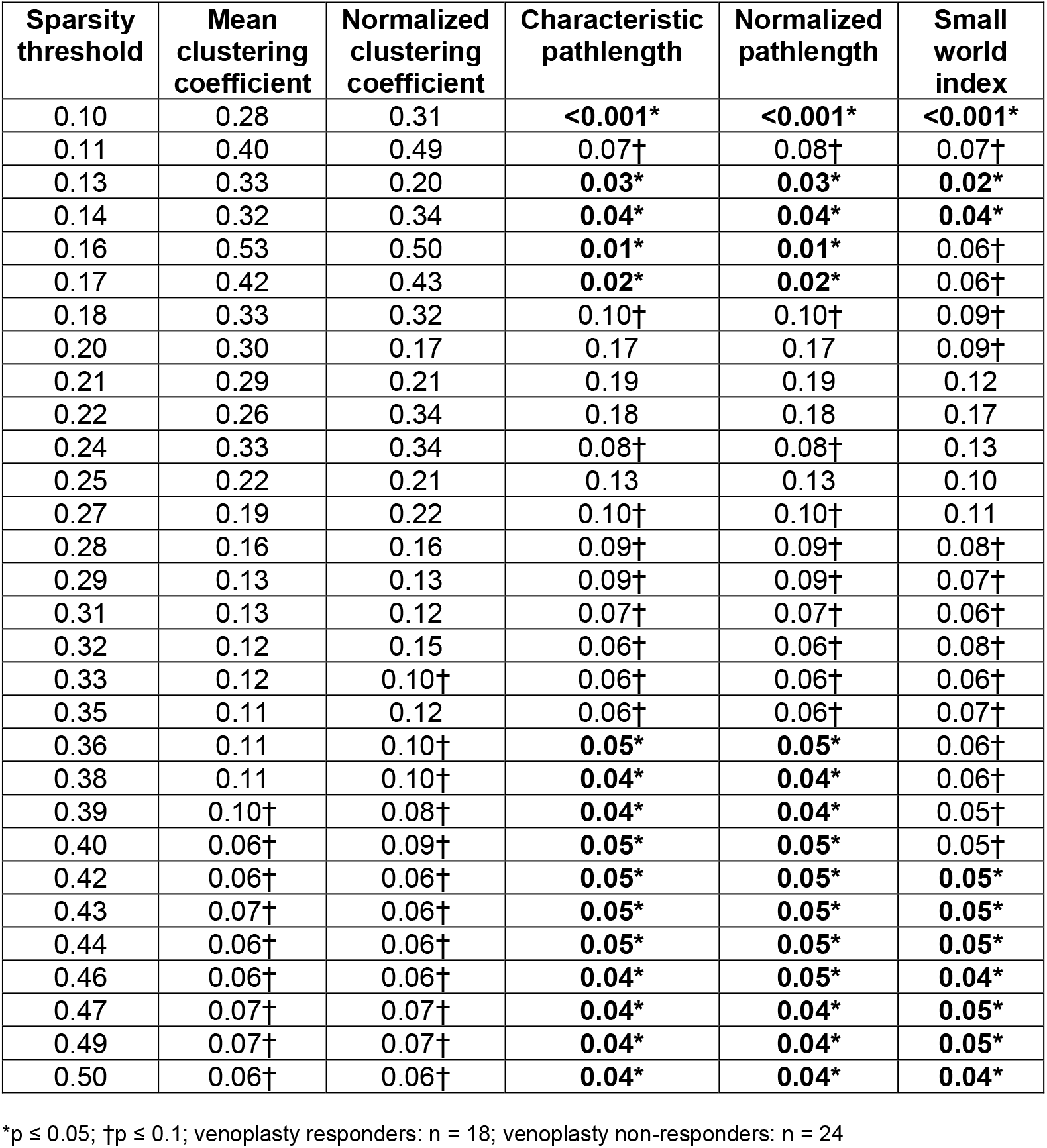
Permutation test p-values for placebo responders versus non-responders including only venoplasty-treated participants at different sparsity thresholds.

